# Flowers respond to pollinator sound within minutes by increasing nectar sugar concentration

**DOI:** 10.1101/507319

**Authors:** Marine Veits, Itzhak Khait, Uri Obolski, Eyal Zinger, Arjan Boonman, Aya Goldshtein, Kfir Saban, Udi Ben-Dor, Paz Estlein, Areej Kabat, Dor Peretz, Ittai Ratzersdorfer, Slava Krylov, Daniel Chamovitz, Yuval Sapir, Yossi Yovel, Lilach Hadany

**Affiliations:** School of Plant Sciences and Food Security, Tel-Aviv University, Tel-Aviv, Israel; School of Zoology, Tel-Aviv University, Tel-Aviv, Israel; School of Mechanical Engineering, Tel-Aviv University, Tel-Aviv, Israel

**Keywords:** pollination, signaling, plant bioacoustics, communication, plant-pollinator interactions, nectar, vibration

## Abstract

Can plants hear? That is, can they sense airborne sounds and respond to them? Here we show that *Oenothera drummondii* flowers, exposed to the playback sound of a flying bee or to synthetic sound-signals at similar frequencies, produced sweeter nectar within 3 minutes, potentially increasing the chances of cross pollination. We found that the flowers vibrated mechanically in response to these sounds, suggesting a plausible mechanism where the flower serves as the plant’s auditory sensory organ. Both the vibration and the nectar response were frequency-specific: the flowers responded to pollinator sounds, but not to higher frequency sound. Our results document for the first time that plants can rapidly respond to pollinator sounds in an ecologically relevant way. Sensitivity of plants to pollinator sound can affect plant-pollinator interactions in a wide range of ways: Plants could allocate their resources more adequately, focusing on the time of pollinator activity; pollinators would then be better rewarded per time unit; flower shape may be selected for its effect on hearing ability, and not only on signaling; and pollinators may evolve to make sounds that the flowers can hear. Finally, our results suggest that plants may be affected by other sounds as well, including antropogenic ones.

## Introduction

Plants’ ability to sense their environment and respond to it is critical for their survival. Plants responses to light (Jiao *et al.* 2007; Chory 2010), mechanical stimulation (De Luca & Vallejo-Marín 2013; Monshausen & Haswell 2013; Appel & Cocroft 2014), and volatile chemicals (Arimura *et al.* 2000; Baldwin *et al.* 2006; Heil & Bueno 2007; Karban *et al.* 2014; Karban 2015) are well documented. However, the ability of plants to sense and respond to airborne sound - one of the most widely used communication modalities in the animal kingdom - has hardly been investigated (Chamovitz 2012; Gagliano *et al.* 2012; Hassanien *et al.* 2014). Recent studies demonstrated slow responses, such as changes in the growth rate of plants, after exposure to artificial acoustic stimuli lasting hours or days (Takahashi *et al.* 1991; Xiujuan *et al.* 2003; Yi *et al.* 2003; Bochu *et al.* 2004; Ghosh *et al.* 2016; Choi *et al.* 2017; Gagliano *et al.* 2017; Ghosh *et al.* 2017; Kim *et al.* 2017; López-Ribera & Vicient 2017; Jung *et al.* 2018). In contrast, to the best of our knowledge, a rapid reaction to airborne sound has never been reported for plants; neither has the biological function of any plant response to airborne sound been identified. In this work we aimed to test rapid plant responses to airborne sound in the context of plant-pollinator interactions.

The great majority (87.5%) of flowering plants rely on animal pollinators for reproduction (Ollerton *et al.* 2011). In these plants, attracting pollinators can increase plant fitness and is achieved using signals such as color, odor, and shape, and by food rewards of nectar and pollen (Willmer 2011). Increased reward quality or quantity can result in longer pollinator visits or in a higher likelihood that a pollinator will visit another flower of the same species in the near future, potentially increasing the flower’s fitness by increasing the chances of pollination and reproduction (Faegri & Van Der Pijl 1979). Producing an enhanced reward can be costly (Pleasants & Chaplin 1983; Southwick 1984; Pyke 1991; Ordano & Ornelas 2005; Ornelas & Lara 2009; Galetto *et al.* 2018) and standing crop of nectar is subject to degradation by microbes (Herrera *et al.* 2008; Vannette *et al.* 2013) as well as to robbery (Irwin *et al.* 2010), including quiet robbers like ants (Galen 1999). Thus, a mechanism for timing the production of enhanced reward to a time when pollinators are likely to be present could be highly beneficial for the plant. Here we suggest that a response of plants to the sound of a pollinator can serve as such a timing mechanism. Specifically, we hypothesize that plants could respond to the sound of a flying pollinator by increasing the reward in a way that would increase the probability of pollination and reproduction by the same or similar pollinators.

The wingbeats of flying pollinators, including insects, birds, and bats, produce sound waves that travel rapidly through air. If plants were able to receive such sounds and react to them rapidly, they could temporarily increase their advertisement and/or reward when pollinators are likely to be present, resulting in improved resource allocation. A possible plant organ that could relay the airborne acoustic signal into a response is the flower itself, especially in flowers with “bowl” shape. If this is the case, we expect that part of the flower (or the entire flower) would vibrate physically in response to the airborne sound of a potential pollinator. We further predict that nectar sugar concentration would increase in response to the sound. None of these predictions have been tested before. To test these predictions we used the beach evening-primrose, *Oenothera drummondii*, whose major pollinators are hawk-moths (at night and early morning), and bees (at dusk and morning) (Eisikowitch & Lazar 1987). We measured petal vibration and nectar sugar concentration in response to sounds. We analyzed the effect of different sound frequencies, including both pollinator recordings and synthetic sounds at similar and different frequencies. We show that pollinator sounds, and synthetic sound signals at similar frequencies, cause vibration of the petals and evoke a rapid response – an increase in the plant’s nectar sugar concentration.

## Materials and methods

### General

We exposed *Oenothera drummondii* plants to different sound playbacks (see below) and measured the concentration of sugar in their nectar. We compared plants’ response to different sounds including pollinator recordings, synthetic sounds in pollinator frequencies and in much higher frequencies, and silence. To determine whether the playback sounds result in physical vibration of the flower petals, we used laser vibrometry. To evaluate pollinator distribution in the field, we performed field observations.

### Experimental setup: Measuring plant nectar response under different treatments

The nectar response was tested in four different experiments (see table S1 for summary): Experiment 1a (n=90 flowers), where the plants were grown outdoors in a natural environment, exposed to natural acoustic conditions, in the summer. The response was tested to the acoustic treatments (see **Sound signals and playbacks** for details): “Silence” – no sound playback, “Low” – playback of a low frequency sound signal with energy between 50-1000 Hz, covering the range of pollinator wingbeat frequencies, and “High” – playback of a high frequency sound signal with energy between 158-160 kHz. This treatment served as a control for the potential effect of the speaker’s electromagnetic field, which was absent in the “Silence” treatment. Experiment 1b (n=167 flowers), where the plants were grown indoors in the summer, and the response was measured to the previous three stimuli plus a “Bee” stimulus - playback of the recordings of a single hovering honeybee with a peak frequency of 200-500Hz; Experiment 2 (n=298 flowers), where the plants were grown indoors in the fall, and response was tested for “Low”, “High”, and “Intermediate” stimuli - playback of a sound signal with energy between 34-35 kHz. To test the role of the flower itself (rather than other parts of the plant exposed to sound) in the response, the “Low” and “High” treatments were also tested for flowers contained in glass jars “Low in Jar” and “High in Jar”; and Experiment 3 (n=112 flowers), where the plants were grown indoors in the spring, and response was tested for “Low” and “High” stimuli.

In each experiment, plants were numbered, randomly assigned to treatments, and tested at a random order, alternating between the different treatments. Different flowers of the same plant were never tested in the same day, nor in the same treatment group. To measure a flower’s response, it was emptied of nectar, and immediately exposed to one of the treatments above. Its newly produced nectar was extracted 3 minutes after the beginning of the treatment (we had to wait three minutes for the amount of nectar accumulated to be measurable by refractormeter). Sugar concentration and nectar volume were quantified before and after the treatment (for details see **Nectar measurements** methods, Fig. S1).

In the jar manipulation, we used 6 identical 1 liter sound proof glass jars, padded with acoustically isolating foam (see Fig. S2). The jar’s ability to block sound was tested by positioning a calibrated microphone (GRAS, 40DP) inside it and playing the “Low” playback from a 10cm distance (as in the experiment). This measurement confirmed that jars reduced sound intensity by 14dB.

### Sound signals and playbacks

In the nectar experiments we used five signals, including bee recordings, three artificial sound stimuli, and silence. The artificial sound stimuli were generated using acoustic software (Avisoft, Saslablite). The “Low” frequency stimulus consisted of a 10s frequency modulated (FM) sound signal sweeping from 1000Hz to 50 Hz, covering the frequency range of the wingbeat of natural pollinators. The “High” frequency stimulus consisted of a 10s frequency modulated sound signal sweeping from 160 to 158 kHz, a frequency that is clearly out of range for pollinator wingbeat. The “Intermediate” frequency stimulus consisted of a 10s frequency modulated sound signal sweeping from 35 to 34 kHz. The “Bee” stimulus was recorded by positioning a calibrated microphone (GRAS, 40DP) and recording an individual honey bee (*Apis mellifera*) from a distance of 10cm. The “Silence” control treatment consisted of no playback.

Acoustic playbacks were performed using a D/A converter (Player 216-2, Avisoft Bioacoustics) at a sampling rate of 500kHz. All signals were recorded using a calibrated microphone before playback to validate their intensity. Playback intensity in the indoor groups “Low”, “Bee”, and “Intermediate” were set to resemble the intensity of a bee hovering 10 cm above the plant, with a peak sound pressure level of ca. 75dB SPL relative to 20µPa at a distance of 10 cm. “Low” playbacks in the outdoor group had a peak pressure of ca. 95dB SPL (relative to 20µPa at 10cm). For control we used either “Silence”, where no sound was played, or the “High” playback which had a weak intensity (ca. 55 dB SPL) but served as an additional ‘Silence’-like condition controlling for the electromagnetic field, absent in the “Silence” control. All playbacks were played continuously for 3 minutes in all treatments, including the silent control. Each playback was played to a group of 5-6 flowers, hovering over each of them with a speaker for a period of 10 seconds each, returning to the first flower at the end. The speakers were moved from plant to plant for 3 minutes at a distance of ca. 10 cm from the nearest flower, mimicking a pollinator hovering around a bush. Thus each flower was exposed to direct sound for 33.8±0.3 seconds on average (we validated that the number of flowers per group had no effect on the significance of the results, see **Results**). Such movement of the speakers was done also in the “Silence” treatment. In both indoor and outdoor experiments, all playbacks were performed indoors: the flowers were brought into a silent room and were treated there.

The vibration experiments were performed with playbacks of a bee (peak energy at 250-500Hz) and a moth (peak energy at ∼100Hz and no energy above 400Hz), and pure tones at the peak frequencies of the signals described above: “Low” (1kHz), “Intermediate” (35kHz), and “High” (160kHz).

### Nectar measurements

Nectar was extracted from all the flowers before treatments using PTFE (Teflon) tubes (external diameter = 0.9 mm, internal diameter = 0.6 mm), followed by disposable 1 µl capillaries for the nectar remaining after emptying by the Teflon tubes. The treatments were applied immediately after extraction. To avoid differences resulting from variation in emptying times we left a capillary inside the first emptied flowers to assure that no new nectar has accumulated. When the last flower was emptied, all capillaries were removed and the treatment (“High”, “Low”, “Bee”, “Intermediate”, or “Silence”) started. Three minutes later, after the treatment ended, nectar was drawn again from all the flowers. Sugar concentration in each flower was measured by calibrated Bellingham-Stanley low-volume nectar Eclipse refractometers (0–50 Brix), where concentration measurements are accurate in volumes as low as 0.2 µL. Three minutes allowed for enough nectar to accumulate in each flower (see Figs S4B, S8B presenting nectar quantities) for the refractometer measurement.

### Monitoring pollinator visitations in the field

In order to assess the pattern of visitation by pollinators in the field, two sets of field observations were done on the Tel Aviv beach. (a) To test whether the presence of a pollinator can indicate the vicinity of additional pollinators, we videoed *Oenothera drummondii* plants during the night. 17 plants were videoed over two nights for four hours after sunset in summer 2017, using IR video cameras (Full Spectrum POV Cam, GhostStop USA, resolution 1920×1080, 30 fps). Cameras were positioned at a distance of 1-1.5 m from the plant. The videos were scrutinized manually using Matlab R2016a and VLC media player 2.2.4. A moth passing within a distance of 1m from a plant was defined as “near the plant”. We then analyzed the distribution of intervals between these events (see results). (b) To estimate the time that a single pollinator spends close to an *Oenothera drummondii* plant, the plants were visually observed during the day, when it was possible to track the same individual over time. 6 plants were observed over four days for three hours in each day. A bee passing within a distance of 20cm from a plant was defined as “near the plant” and the time it spent within this distance was estimated.

### Measuring petal vibration using laser vibrometry

To determine whether the playback sounds result in physical vibration of the flower petals, we used laser vibrometry. This method allows measuring minute physical vibrations through doppler shifts of a laser beam reflected from a vibrating surface. To this end, the flowers were positioned on a wafer prober (Karl Suss PSM6, Mitutoyo FS70L-S microscope) and operated in ambient air. The motion of the petals was registered using a laser Doppler vibrometer (Polytec LDV, OFV-5000 controller). The vibrometer was operated in the velocity acquisition mode using VD-02 Velocity Output Decoder, (up to 1.5 MHz bandwidth). The laser beam was focused on the base of the petal (see Fig. S3) using the x5 long working distance lens of the microscope.

Signals from the LDV were fed into the oscilloscope KEYSIGHT DSOX2004A (70 MHz, 1 Mpts memory). We compared flower vibrations in response to different playback frequencies and in the absence of playback (“Silence”) in a paired experimental design (within the same plant). To validate that the presence of petals was crucial to the vibration we also compared petal vibration in intact flowers to vibration in intact petal of flowers where some of the petals were removed (see Figs. S3).

To measure the actual vibration amplitude, we subdivide the measured velocity by 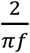, where *f* is the frequency of the oscillation. We used vibration models of objects with similar shapes (both a beam and a circular thin plate (Blevins & Plunkett 1980) to estimate the flower’s resonance vibration frequency. The resonance frequency of an object is dictated by the material properties, geometry and boundary conditions. For flower size of ∼6 cm and thickness of ∼0.4 mm we estimated a fundamental mode frequency to be in the range of 100-500 Hz. A measured density of ∼230 kg/m3 and the Young’s modulus of ∼1 MPa adopted from (Watanabe & Ziegler 2013) were used in calculations.

### Plants and growth conditions

*Oenothera drummondii* plants were propagated from grafts of plants taken from Bet-Yanai coast, Israel. In all experiments, irrespective of the plants growth conditions, the response of the plants to sound playback was tested indoors, in a quiet room. For experiment 1a (“outdoor, summer 2014”) 200 plants were placed in 3 liter pots and grown in the Tel Aviv University Botanical Gardens in an outdoor setting. Flower buds were covered with nets a day before the experiment, to avoid pollination and nectar withdrawal by pollinators. For experiment 1b (indoor, summer 2015) 100 plants were placed in 0.5 liter pots, for experiment 2 (indoor, fall 2016) 400 plants were placed in 1.1 liter pots, and for experiment 3 (indoor, spring 2016) 200 plants were placed in 0.5 liter pots. Experiments 1b, 2 and 3 used indoor-grown plants only, as outdoor plants flower only in the summer. For all indoor experiments the plants were grown in a controlled growth room, at 27-28 degrees centigrade, with 16 hours of artificial daylight, about 1 month prior to the beginning of the experiment. Altogether, more than 650 flowers from these 900 plants were used in the nectar experiments, and another ∼200 flowers in the laser experiments (taken from the plants of experiments 2 and 3). In each experiment, only plants of the same age and season were tested. See Table S1 for summary of the experiments.

### Statistical analysis

Experiment 1: We performed a two-way ANOVA on log sugar concentration, including the treatment (“Silence”, “High”, “Low”, or “Bee”) and group (“indoor” or “outdoor”) variables. The group variable was not found to have a significant effect (P = 0.793). Therefore, data from both groups were combined and the sugar concentration and nectar volume between different treatments was compared. Shapiro-Wilks test concluded significant deviation from normality (P<0.05) in some of the cases (nectar volume data), so Wilcoxon rank-sum was used for comparison. Within groups, the reported p values were adjusted for multiple comparisons using the Holm–Bonferroni method.). Experiment 2: nectar traits (sugar concentration and nectar volume) under different treatments were compared using Wilcoxon rank-sum. To test the effect of hydration status (days since watering), number of flowers in group, and time of day on our results we used analysis of variance (ANOVA) model. We used log (sugar concentration) as the dependent variable, and hydration status (or number of flowers in group or time of day), treatment group, and their interaction as predictors. Post-hoc p-values were calculated using a Tukey HSD test. Constant variance assumption was corroborated using Levene’s Test. All petal vibration levels were compared using paired Wilcoxon test (comparing vibration levels of the same flower under different treatments or petal removal). Pollinator distribution in the field was compared using paired Wilcoxon test as well.

## Results

We found that *Oenothera drummondii* flowers produced nectar with significantly increased sugar concentration (Wilcoxon p<0.01, Fig. 1A; and see methods) after exposure to the playback of the natural sound of bee wingbeats, as well as in response to artificial sounds containing similar frequencies (the “Bee” and “Low” treatments, Fig. 1B middle and right), in comparison with flowers exposed to either high frequency sounds (the “High” treatment, Fig. 1B left) or no sound at all (the “Silence” treatment). The average sugar concentration was 20% higher in flowers exposed to pollinator-like frequencies (“Bee” and “Low” sound signals), in comparison with flowers exposed to “Silence” or “High”, while no difference was observed between flowers exposed to “High” frequencies and flowers exposed to the “Silence” treatment. No difference in sugar concentration was observed between experimental groups before the treatment, and the volume of the nectar produced by the flowers did not change significantly in the “Bee” and “Low” treatments (Fig. S4), showing that the increase in sugar concentration in these groups could not be attributed to a decrease in water volume. Analyzing the data using Student’s t-test of log-transformed data resulted in similar significant results (P < 0.001 for each of the comparisons between treatment (“Low” and “Bee”) and control (“High” and “Silence”).

**Fig 1.**
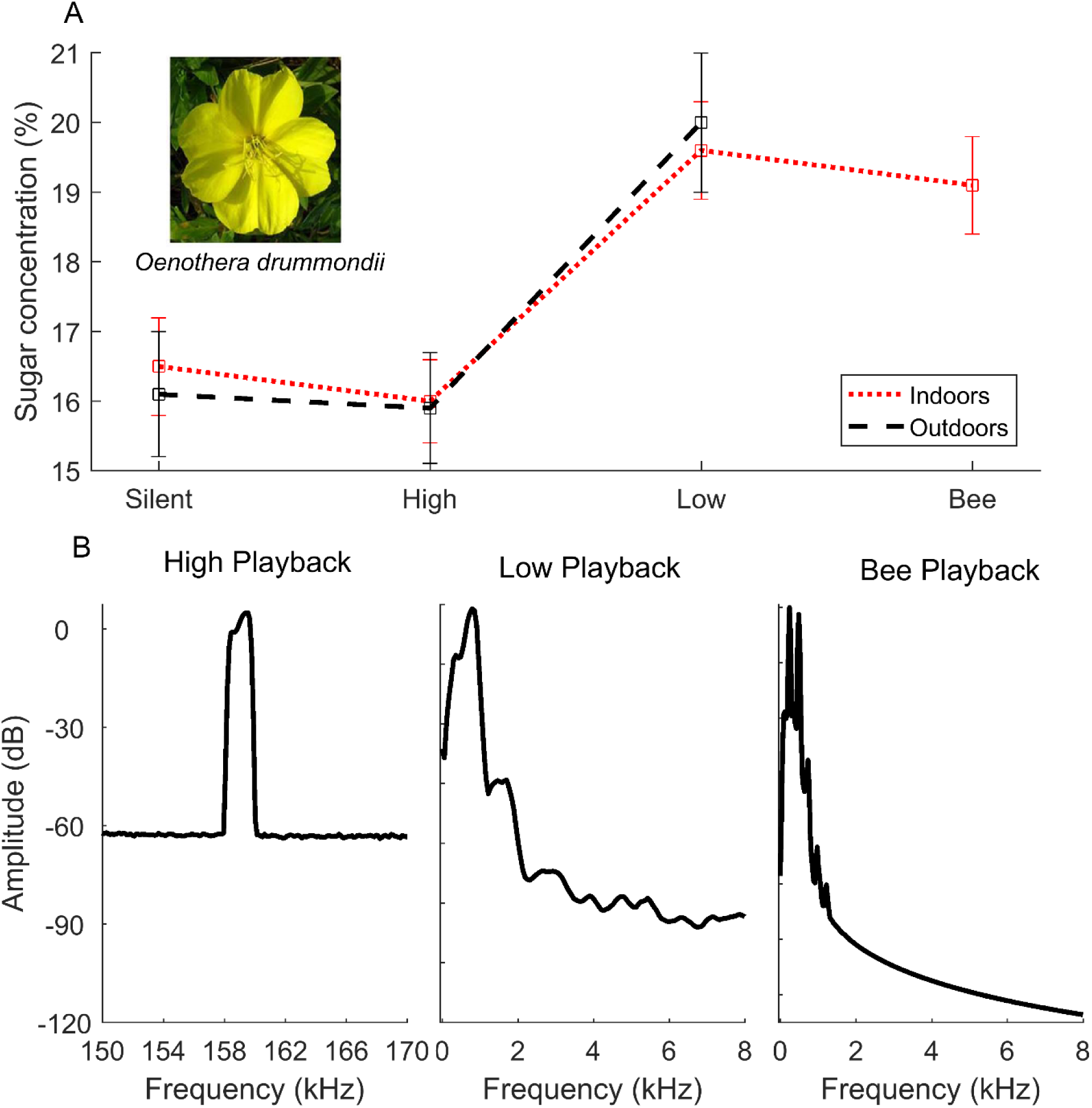
Flowers respond rapidly to pollinator sounds by producing sweeter nectar. A. Mean sugar concentration under the different treatments in outdoor (dashed black) and indoor (dotted red) experiments. Mean sugar concentration across both indoor and outdoor groups differed significantly (P<0.01) between flowers exposed to frequencies below 1kHz (sugar concentration 19.8% ± 0.6, n=72 and 19.1% ± 0.7, n=42 for “Low” and “Bee” after 3 minutes, respectively), compared to flowers exposed to “Silence” or “High” frequency sound (16.3% ± 0.5, n=71, and 16.0% ± 0.4, n=72, respectively). Insert shows a flower of *Oenothera drummondii*. B. Spectra (frequency content) of the playback signals used in the experiment. Both “Bee” and “Low” signals contain most energy below 1000Hz, while the “High” control peaked at ca. 159,000Hz.

To verify the potential advantage of increasing sugar concentration within minutes after a pollinator’s sound, we video-monitored the distribution of pollinators near *Oenothera drummondii* flowers in the field over two nights. We found that one pollinator flying in the vicinity of the plant – and producing sound in the process – is a strong indication that another or same individual may be in the plant vicinity within a few minutes. Specifically, a pollinator was >9 times more common near the plant if a pollinator was near the plant in the preceding 6 minutes, than if no pollinator was around in the preceding 6 minutes (see Fig. S5 and Monitoring pollinator visitations methods). A response of the plant within minutes to the sound of a nearby flying pollinator could thus serve to better reward another pollinator in the vicinity (or possibly the same individual). We further quantified the time pollinators tend to stay nearby *Oenothera drummondii* flowers in the field (Methods). Two species of bees were observed around the flowers, and the observed “buzzing times” (near the flower) were 27.8 ± 7.7 for honey bees (n=44), and 38.9 ± 11.8 for carpenter bees (n=23), see Fig. S6. In reality, plants may of course be exposed to longer sound stimuli due to multiple bee passes one after the other. Notably, as our playback lasted 3 minutes and we had 6 plants at each session, each plant was exposed to 30 seconds of direct playback, on average.

To determine whether the sound waves emitted by a pollinator result in physical vibrations of the flower, we used laser vibrometry (see methods). *Oenothera drummondii* flowers vibrated mechanically in response to the airborne sounds of a bee or a moth recording (Fig. 2A, and S7 for moth sound spectra), oscillating in velocities that have already been shown to elicit a defense response in a plant that was mechanically moved in such velocities (Appel & Cocroft 2014). The amplitude of the mechanical vibrations (which reached 0.1mm) depended on the presence of intact petals, and significantly decreased upon removal of petals (Fig. 2B, P<0.0005, see Fig S2 for details), suggesting that the petals either directly receive, or serve to enhance the received signal.

**Fig 2.**
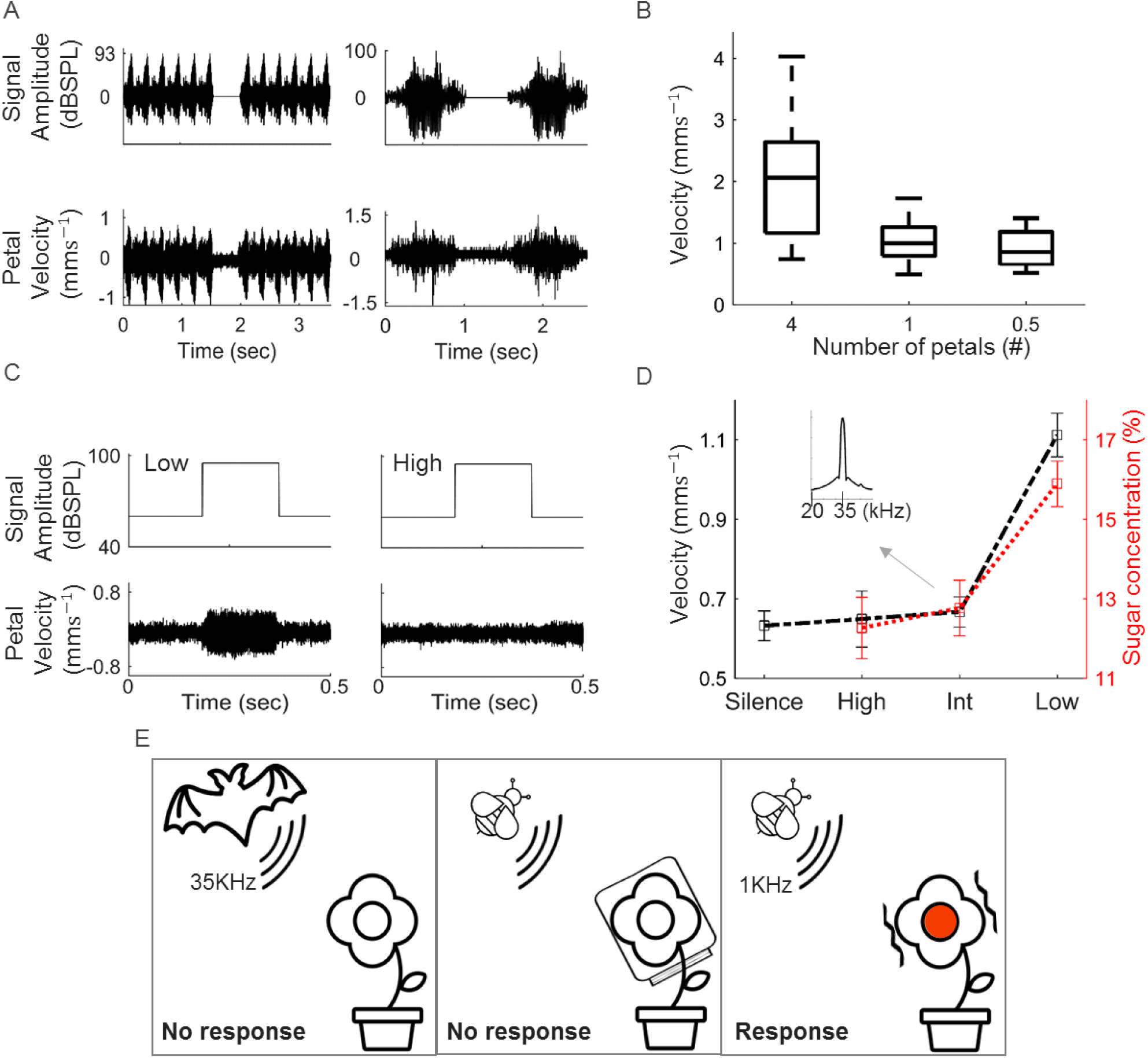
A. Flowers vibrate mechanically in response to airborne sound of a pollinator. Top: Left - time signal of a honey bee sound signal (airborne signal recorded using a microphone). Right - time signal of a flying *Plodia interpunctella* male moth (the signal’s spectrum peaks at ∼100Hz, see Fig. S7). Bottom: Mechanical vibration recorded in an *Oenothera drummondii* flower in response to the playback of the bee (left) and moth (right) sound signals. B. Vibration amplitude in response to the bee signal depended on the presence of petals: a significantly stronger vibration was recorded when all 4 petals were intact in comparison to when flowers were trimmed and had only 1 or 0.5 petals (paired Wilcoxon, P<0.0005 for the comparison between 4 and 1 petal and P<0.005 for the comparison between 4 and 0.5 petals). C. Flowers vibrated in response to playback of low frequencies around 1kHz (left) while they did not vibrate above background noise to playbacks at higher frequencies of ∼35kHz (right). Top: the time that the playback was ‘on’. Bottom: Vibration time signals of the flowers. D. Frequency specificity in both vibration and sugar concentration response. The flowers vibrated (dashed black) significantly more than background noise in response to sound signals in low frequencies around 1kHz (paired Wilcoxon p<0.0001, n=21) but not in response to high frequencies around 160kHz (p>0.6, n=23) or to intermediate frequencies around 35kHz (p>0.9, n=21); The flowers also increased sugar concentration (dotted red line) in response to “Low” signals significantly more than in response to the “Intermediate” signal presented in the inset (p<0.002), or to the “High” signal serving as control (p<0.0001). (Sugar concentration 15.9% ± 0.57, n=81, 12.8% ± 0.7, n=49, and 12.3% ±0.77, n=51, for Low, High and Intermediate, respectively). Inset shows the spectrum of the “Intermediate” playback signal used in the nectar experiment. E. Summary of experimental results. Flowers vibrate in response to airborne sound at pollinator’s frequency range, and increase nectar sugar concentration (right panel). Glass covered flowers do not respond (middle), suggesting that the flower serves as the plant’s “ear”. The flowers response is frequency specific, and they do not vibrate or respond to frequencies around 35 kHz (left).

To test the frequency-specificity of the response we performed another indoor experiment (experiment 2, in the fall) in which we repeated the use of the previous sound stimuli (Low and High) and introduced another “Intermediate” sound signal with a peak frequency of 35kHz (240 new flowers were used in this experiment, in the fall, see Table S1). The flowers showed frequency specificity, both functionally and mechanically: they vibrated significantly (paired Wilcoxon p<0.0001, n=21) in response to sound signals of the “Low” signal, 1kHz, but not in response to the peak frequency of an “Intermediate” signal, 35kHz (p>0.9, n=23), or the “High” signal, 160kHz (p>0.9, n=21, see Fig. 2C and 2D black line). Similarly, the flowers increased sugar concentration in response to “Low” sound signals significantly (p<0.002, 2D red dotted line) in comparison to the “Intermediate” or “High” treated flowers. “Low” sounds resulted in significantly higher sugar concentration than all other treatments (High, Intermediate, High in jar, Low in jar) also when accounting for hydration status, number of flowers in group, or the time of day (p<0.03). Differences between the four other treatment groups were not significant. The ratios of post-treatment to pre-treatment concentration and vibration, per plant, revealed an identical pattern: the ratios were significantly higher (p<0.002 for concentration ratio, see Fig. S8, p<e-07 for vibration ratio) in plants exposed to “low” sounds in comparison with plants exposed to “high” or “intermediate” sounds (Table S2). In another experiment (experiment 3, n=112 flowers, in spring) where only “Low” and “High” stimuli were tested, there was again significant increase in the sugar concentration in response to the “Low” stimuli (see Table S3). The plants showed different flowering phenotype in different seasons probably reflecting the season experienced before entering the growth room: summer plants (Experiment 1) had larger flowers with higher sugar concentration before treatment in comparison with either fall plants (Experiment 2) or spring (see Table S4). Regardless, the major pattern – an increase in nectar sugar concentration in response to pollinator sound playbacks – was highly significant in all seasons (Fig. 2D, 1A, Table S3).

Finally, to validate the importance of the flower itself as the organ responsible for sound reception, we ran another experiment. When the flowers (but not the stem or leaves) were covered with glass jars that blocked sound (see Fig. S2), then the “Low” playback had no effect on the sugar concentration: For flowers enclosed in jars, there was no significant difference between exposure to “Low” treatment and exposure to “High” (p>0.64, for n=58 and 59 flowers, respectively), and none of these groups differed significantly from the no jar “High” treatment, that served as a control (p>0.49, n=49 flowers, see Table S2).

## Discussion

We found that plants respond rapidly to specific airborne sound frequencies (Fig. 1, 2D) in a way that could potentially increase their chances of pollination, and that flowers can serve as sound sensing organs (Fig. 2). Consistent results were obtained in four independent experiments (Table S1) with over 650 flowers in total. The flowers responded similarly to bee wingbeat sounds and to artificial sound-waves that were similar in their frequency spectrum but differed greatly in their temporal pattern, suggesting that the frequency of the sound is sufficient to elicit a response. The flowers responded rapidly, within 3 minutes. The concentration of sugar in the nectar produced following the exposure to sound increased by ∼20% on average.

Bees have been shown to be capable of perceiving differences in sugar concentration, as small as 1-3% (Afik *et al.* 2006; Whitney *et al.* 2008). Thus, even if the new sugar-rich nectar is diluted by lower concentration nectar already present in the flower, the bees would be able to detect the difference in many cases. This is already true 3 minutes after first sound emission, and the absolute sugar concentration could further increase if the plant’s response continues. Increased sugar concentration can enhance the learning process of the pollinators, and facilitate pollinator constancy – the tendency to visit flowers from the same species (Cnaani *et al.* 2006) – thus increasing the effectiveness of pollination. Enhanced reward can also increase visit duration, further enhancing pollination efficiency (Manetas & Petropoulou 2000; Brandenburg *et al.* 2012). This is not without caveats: too high sugar concentration could result in too viscous nectar for some pollinators, but the values measured here are below the optimum for both bees and moths (Josens & Farina 2001; Krenn 2010; Kim *et al.* 2011), suggesting that the polinators can benefit from the increased concentration. It may also result in a higher number of flowers visited per plant, possibly leading to geitonogamous selfing (Klinkhamer & de Jong 1993; Hodges 1995; Dafni *et al.* 2005). Yet, if only part of the flowers in the plant carry enhanced rewards – e.g., due to depletion – then the response could result in increased variation in nectar standing crop within the plant, encouraging the pollinators to move to the next plant and facilitating outcrossing (Ott *et al.* 1985; Biernaskie & Cartar 2004; Pyke 2016).

A response within 3 minutes is advantageous when pollinators move between nearby flowers, or when the presence of one pollinator is a good predictor of other nearby pollinators, such as in bees (Goulson 1999; Slaa *et al.* 2003) and in moths according to our field observations (Fig. S5). Such a response would allow the plant to identify the beginning and intensity of pollinator activity which can differ from day to day due to various factors such as weather conditions (Corbet *et al.* 1993). The plant could then switch to an increased sugar production mode, in order to reward the first actual visitors. Rapidly increasing nectar sugar concentration would be advantageous also in the case of a sporadic pollinator remaining in the area of the plant for a long time. Note that in a plant like the evening primrose, characterized by multiple flowers (dozens of flowers in a mature bush), the response to the sound of a nearby pollinator could be beneficial even if the pollinator avoids visiting the specific flowers that had recently been visited (Giurfa & Núñez 1992; Goulson *et al.* 1998), since it can still visit other flowers of the same plant. Other pollinators actually prefer occupied or recently-occupied food sources (Schmidt *et al.* 2003; Kawaguchi *et al.* 2006; Lihoreau *et al.* 2016), and might especially benefit from enhanced refilling.

The plants responded to specific sound frequencies characteristic of pollinators’ wingbeat (Figs. 1, 2). How could such specificity be attained? We estimated the resonance frequency of the evening primrose petal to be a few hundred Hz based on vibration models developed for objects with similar shapes (Blevins & Plunkett 1980). This is close to the sound frequencies typically generated by bee and moth wingbeat. Furthermore, the flower should vibrate mostly around the resonance frequency, and vibrate less in response to higher or lower frequencies. This could explain how the flower increased sugar concentration in its nectar only in response to low frequencies. This frequency specificity might also explain how the flower filters wind-induced vibrations, which are typically at lower frequencies (Appel & Cocroft 2014).

The current work is a first step in a new field, and can be extended in several ways. First, the response to sound can be further studied in the wild, on the background of other natural sounds. Second, all our nectar measurements were performed by first emptying the flower and then measuring refilled nectar. Testing the response to sound without prior manipulation will be more realistic (Corbet 2003), but would require large sample sizes due to the high variation in the nectar standing crop present in the model species. Third, the actual functionality of the response has yet to be tested – i.e., do pollinators indeed prefer plants exposed to sound, and to what extent? Fourth, we tested the response to sound in a single plant species. Additional species might reveal different responses according to their specific ecologies (e.g., bat pollinated plants may respond to different frequencies).

The petal vibrations that we measure could be picked up by mechanoreceptors, which are common in plants (Monshausen & Gilroy 2009), and have been shown to respond to vibrations with similar amplitudes (Appel & Cocroft 2014). We hypothesize that the flower serves as the plant’s external “ear” in terms of receiving sound pressure. We posit that the petals of other flowering species could have evolved to detect sound, similar to our findings in *Oenothera drummondii*. The resonance frequency of a flower will be dictated by its mechanical parameters: size, shape and density, which could be under natural selection. If plant responses to airborne acoustic signals are indeed adaptive in the context of pollination, we expect plants with “noisy” pollinators – such as bees, moths, and birds – to have evolved large ear-like flowers with proper mechanical parameters making them sensitive to the sounds of their pollinators.

Much is known about the response of pollinators to plant signaling from a distance (Patiny 2011; Schaefer & Ruxton 2011). In contrast, the response of plants to pollinators from a distance has never been demonstrated. The implications of such a response to the ecological system might be far reaching, since pollination is critical for the survival of many plant species, including many agriculturally important crops (Kremen *et al.* 2002; Faegri & Van der Pijl 2013). Plant response to sound could allow bi-directional feedback between pollinators and plants, which can improve the synchronization between them, lowering nectar waste, and potentially improving the efficiency of pollination in changing environments. These advantages can be diminished in very noisy environments, suggesting possible sensitivity of pollination to external noises, including antropogenic ones. Finally, plants’ ability to hear has implications way beyond pollination: plants could potentially hear and respond to herbivores, other animals, the elements, and possibly other plants.

## Author Contributions

LH, YY, YS, and DAC designed the research. MV, IK, UBD, PE, AB, AK, DP, IR, AG, KS, and EZ performed the experiments. IK, UO, MV, YY, and LH analyzed the data. YY LH and SK supervised the acoustic experiments. YS and LH supervised the nectar experiments. LH, YY, and YS contributed equally to the study. All authors discussed the results and took part in writing the manuscript.

## Acknowledgements

We thank Prof. Dan Eisikowitch, Prof. Amram Eshel, Dr. Iftach Vaknin, and Dr. Yael Mendelik for contributing nectar measuring equipment; Stella Lulinski for help with the laser experiments; Stav Hen, Dorin Cohn, Oren Rabinowitz, and Ran Perelman for help with the nectar experiments; Prof. Nir Ohad, Dr. Tuvik Beker, and Prof. Judith Berman for comments on the manuscript. The research has been supported in part by Bikura 2308/16 (LH, YS, YY), Bikura 2064/18 (LH, YS, YY), ISF 1568/13 (LH), by the Smaller Winnikow fellowship (MV), and by the Manna Center Program for Food Safety and Security fellowships (IK, UO).

