## Supplementary Data for "Flowers respond to pollinator sound within minutes by increasing nectar sugar concentration"

### Supplementary figures and tables

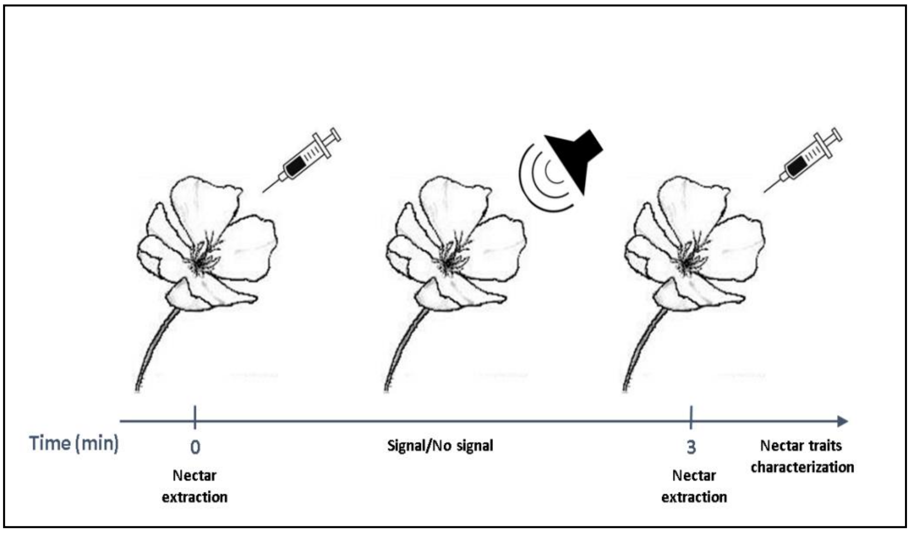


**Figure S1.** Course of the nectar experiments. First all the nectar was extracted from the flower (time 0). Then the flower was exposed to the relevant signal (‘High’, ‘Low’, ‘Intermediate’ or ‘Bee’ playback). After 3 minutes the new nectar was extracted again and its traits were characterized.

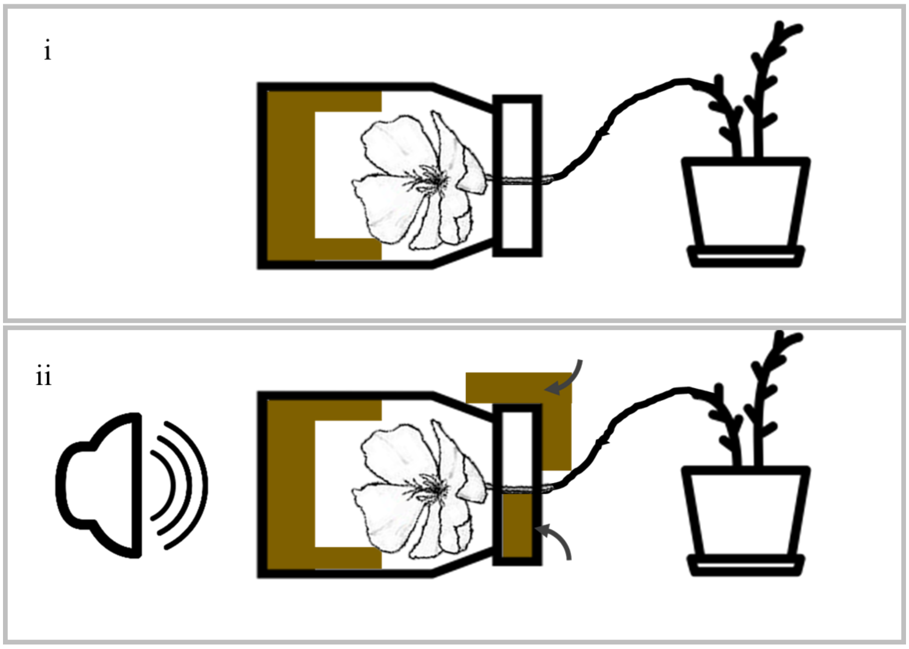


**Figure S2.** Course of jar experiment. First the flower was inserted into a 1-liter sound proof glass jars, padded with acoustically isolating foam (panel i). The opening of the jar was also blocked with 2 pieces of acoustically isolating foam before the signal was played (panel ii).

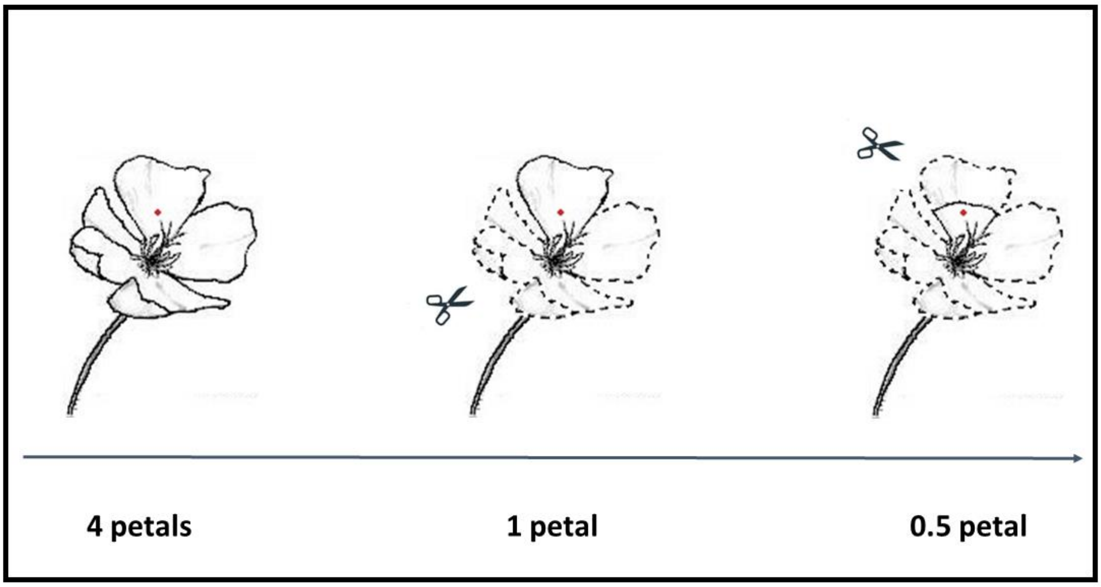


**Figure S3. Course of petal removal in the petal vibration tests**. The vibration of the same flower was measured 3 times while the bee signal was played – first with all of its 4 petals (left), then 3 petals were removed, leaving one intact petal (middle) and finally the last petal was trimmed at around half it's length (right). The laser was focused on the red dot. This process was performed on all the flowers in the experiment.

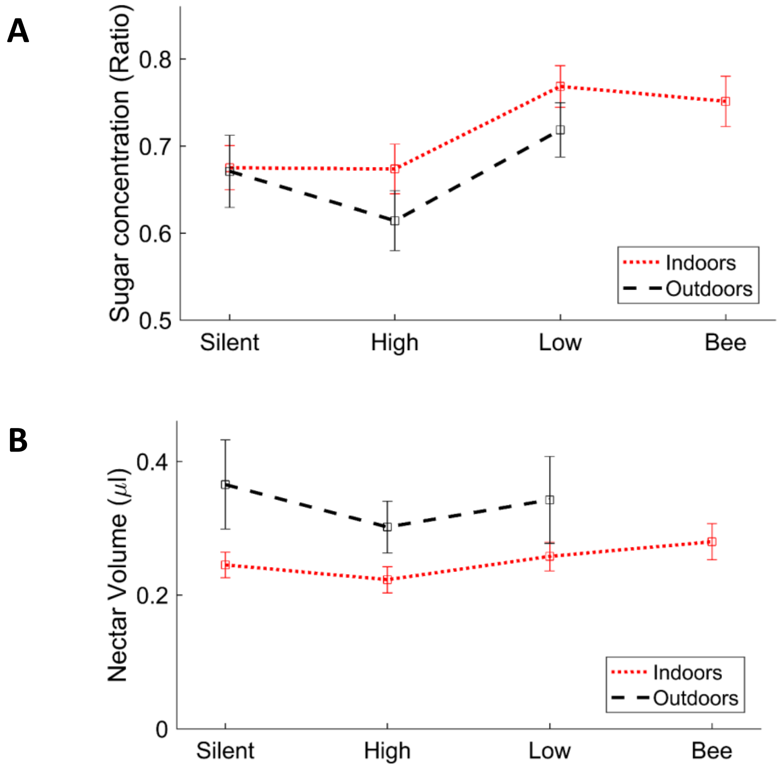


**Figure S4.** A. **Mean sugar concentration ratio, experiment 1** ("sugar concentration after stimuli"/"sugar concentration before stimuli") under the different treatments in outdoor (dotted red) and indoor (dashed black) experiments. Mean sugar concentration ratio across both indoor and outdoor groups differed significantly (P<0.02), between flowers exposed to "Low" frequency sound (sugar concentration ratio 0.75 ± 0.019, n=42), compared to flowers exposed to "Silence" or "High" frequency sound (0.67 ± 0.022, n=71 and 0.65 ± 0.022, n=72, respectively). Mean sugar concentration ratio differed significantly (P<0.02) between flowers exposed to "Bee" frequency sound (sugar concentration ratio 0.75 ± 0.029), compared to flowers exposed to "High" frequency sound (0.65 ± 0.022). However, it did not differ significantly (P=0.07) between flowers exposed to "Bee" frequency sound (sugar concentration ratio 0.75 ± 0.029), compared to flowers exposed to "Silence" treatment (0.67 ± 0.022). No significant difference in sugar concentration was observed between any two groups before the treatment. B. Mean nectar volume under the different treatments in outdoor (dotted red) and indoor (dashed black) experiments. Mean nectar volume did not differ significantly (P>0.25) between flowers exposed to frequencies below 1kHz (nectar volume 0.29 μl ± 0.03 and 0.28 μl ± 0.02 for "Low" and "Bee" after 3 minutes, respectively), compared to flowers exposed to "Silence" or "High" frequency sound (0.29 μl ± 0.03 and 0.25 μl ± 0.02, respectively).

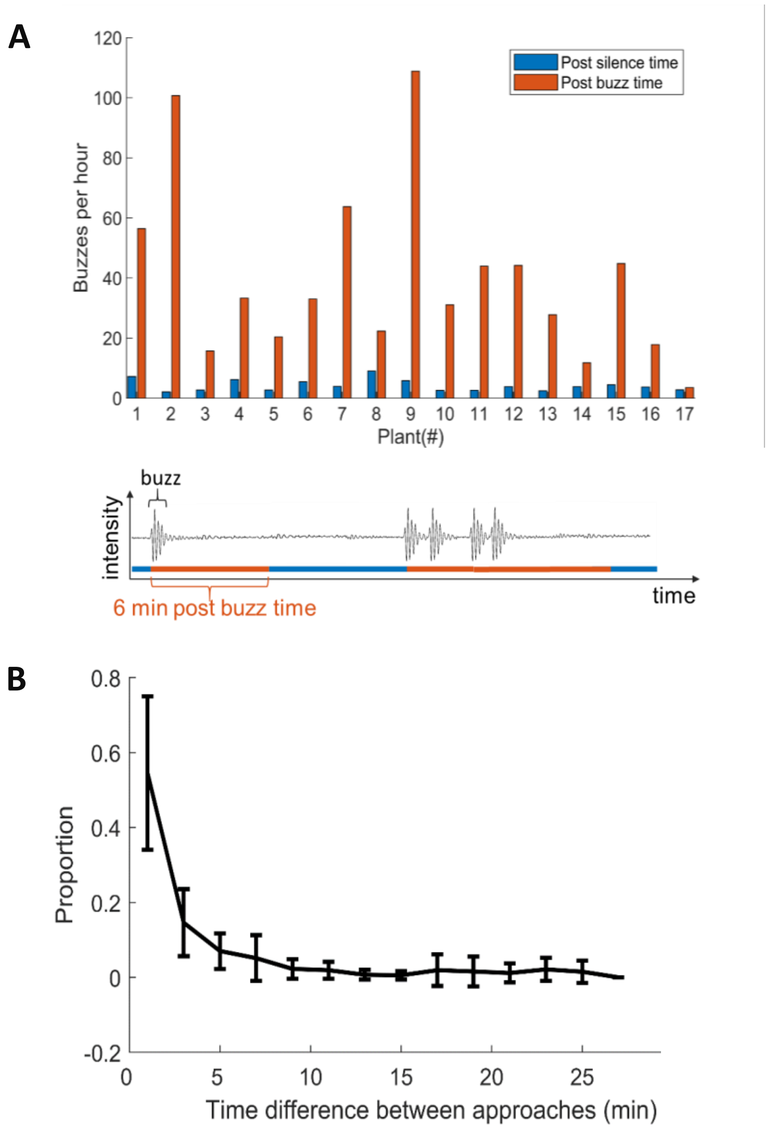


**Figure S5. The presence of a pollinator near the plant is a good predictor for another pollinator within minutes. A.** Top: The mean rate of pollinators flying next to a plant during *post buzz times* – time preceded by a pollinator flying near the plant in the last 6 minutes (red) – was significantly higher (P <0.001) than the rate at all other times (*post silence time*, blue). 17 plants were recorded and the data for each plant is shown separately (X-axis). Bottom: Illustration of the calculation of *post buzz time.* We divided the entire time into *post-buzz* periods (red) and *post silence* periods (blue). *Post buzz periods* were defined as 6 minute windows after a pollinator's approach of the plant (depicted by a pulse in the image). If another pollinator arrived during these 6 minutes (see the third buzz in the illustration), "*post buzz time*" extends until 6 minutes after the last buzz. *Post silence time* includes all the time not included in *post buzz time*, namely times preceded by 6 or more minutes without a nearby pollinator. We calculated the numbers of pollinators flying near the plant during *post buzz (or post-silence)* time, and normalized by the total length of *post buzz (or post silence)* time respectively. **B.** Distribution of the intervals between buzzes.

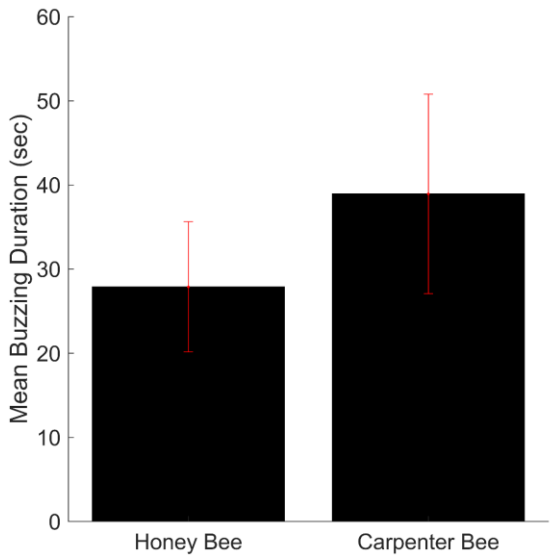


**Figure S6.** Buzzing duration of honey and carpenter bees in the flower area. The buzzing duration was calculated for each bee and was defined as the time that the bee spent in a radius of 10cm from the plant. $N_{Honey-bee} = 44, N_{Carpenter-bee} = 23$ .

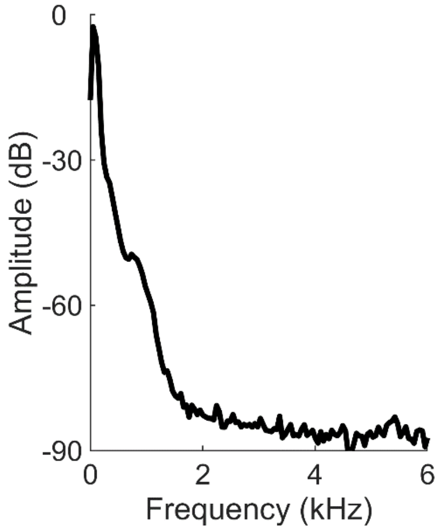


**Figure S7.** Spectra of the moth playback signal used in the experiment. Like the bee and the low frequency signals, the moth signal contains most energy below 1000Hz.

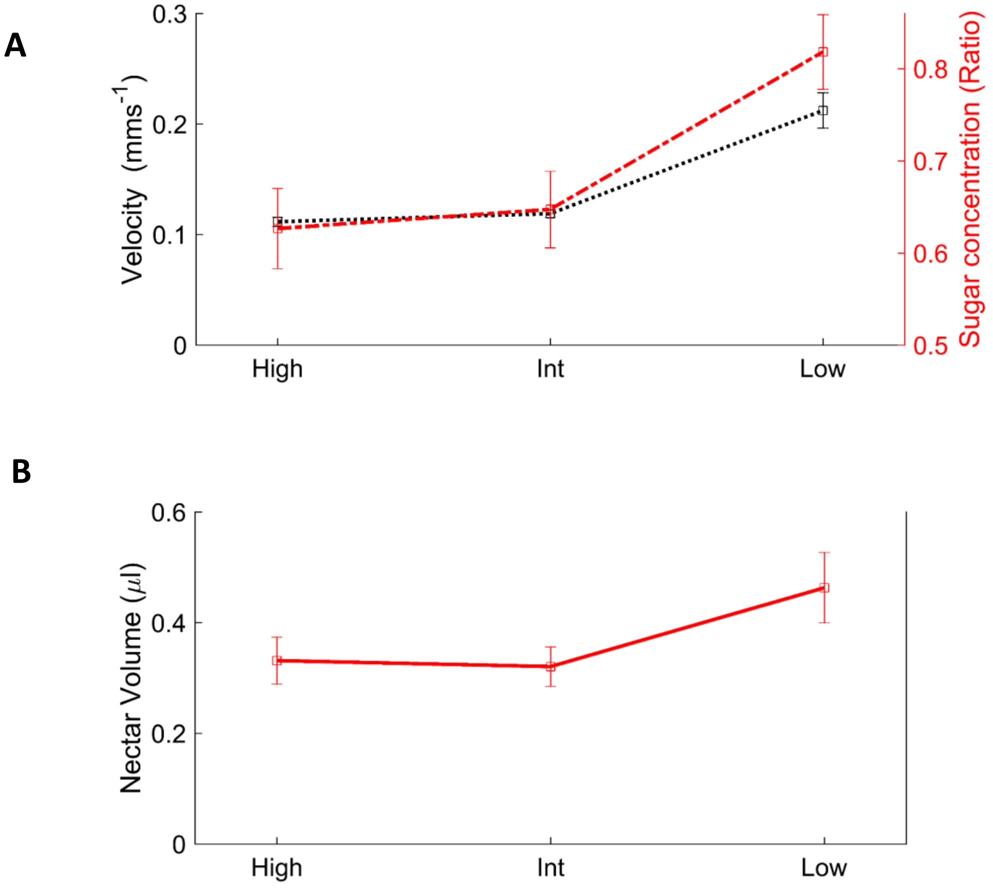


**Figure S8.** A. **Mean sugar concentration ratio, experiment 2** ("sugar concentration after stimuli"/"sugar concentration before stimuli", dotted red) and mean petal vibration (velocity in $mms^{-1}$, dashed black) under the different treatments in Experiment 2. Mean sugar concentration ratio differed significantly (P<0.002) between flowers exposed to “Low” frequency sound (sugar concentration ratio 0.82 ± 0.04, n=81), compared to flowers exposed to “Intermediate” or “High” frequency sound (0.65 ± 0.04, and 0.62 ± 0.04, respectively, n=51 and 49 respectively). Mean petal vibration differed significantly (P<6.35e-08) between flowers exposed to “Low” frequency sound (velocity 0.20 $mms^{-1}$ ± 0.014, n=20), compared to flowers exposed to “Intermediate” or “High” frequency sound (0.118 $mms^{-1}$ ± 0.003 and 0.111 $mms^{-1}$ ± 0.004, respectively, n=20 each). B. **Mean nectar volume** under the different treatments. Mean nectar volume did not differ significantly (P>0.29) between flowers exposed to "Low" frequency sound (nectar volume 0.46 μl ± 0.03), compared with flowers exposed to “Intermediate” or “High” frequency sound (0.32 μl ± 0.03 and 0.33 μl ± 0.04, respectively).

| Exp. | Name | Plants (*Oenothera drummondii* seedlings) | N (flowers per exp.) | Growth conditions | Treatment stimuli (frequency range) | Control stimuli (frequency range) | Exp. location | Presented at |
|---|---|---|---|---|---|---|---|---|
| 1a | Outdoor – summer 2014 | 200 | 90 | outdoors at the Botanical Garden, in 3 liter pots | 1. "Low" – 50-1,000 Hz | 1. "Silence" – silent speakers moving around the flower<br>2. "High" – 158-160 kHz | In a quiet room | Figure 1A, Figure S4 |
| 1b | Indoor – summer 2015 | 100 | 167 | controlled growth room, in 0.5 liter pots | 1. "Low" – 50-1,000 Hz<br>2. "Bee" – 200-500 Hz | 1. "Silence" – silent speakers moving around the flower<br>2. "High" – 158-160 kHz | In a quiet room | Figure 1A, Figure S4 |
| 2 | Indoor – fall 2016 (double-blind, mechanism and specificity) | 400 | 298 | controlled growth room, in 1.1 liter pots | 1. "Low" – 50-1,000 Hz<br>2. "Intermediate" – 34-35 kHz<br>3. "Low in Jar" – 50-1,000 Hz<br>4. "High in jar" – 158-160 kHz | 1. "High" – 158-160 kHz | In a quiet room | Figure 2D, Figure S8, Table S2 |
| 3 | Indoor – spring 2016 (double-blind) | 200 | 112 | controlled growth room, in 0.5 liter pots | 1. "Low" – 50-1,000 Hz | 1. "High" – 158-160 kHz | In a quiet room | Table S3 |

**Table S1: Nectar experiments summary.**

**A**

| Treatment | N | Sugar concentration (%) | | | | Sugar concentration ratio | | | |
|---|---|---|---|---|---|---|---|---|---|
| | | mean | SE | P (Wilcoxon rank-sum) | | mean | SE | P (Wilcoxon rank-sum) | |
| | | | | High | Low | | | High | Low |
| High | 49 | 12.27 | 0.77 | | 0.0004 | 0.62 | 0.04 | | 0.003 |
| High in jar | 59 | 13.14 | 0.55 | 0.49 (0.12) | 0.005 | 0.65 | 0.02 | 0.80 (0.20) | 0.006 |
| Low in jar | 58 | 12.81 | 0.62 | 0.92 (0.30) | 0.003 | 0.67 | 0.04 | 0.96 (0.32) | 0.01 |
| Int. | 51 | 12.77 | 0.70 | 0.95 (0.47) | 0.005 | 0.64 | 0.04 | 1 (0.63) | 0.007 |
| Low | 81 | 15.90 | 0.57 | 0.0004 | | 0.81 | 0.04 | 0.003 | |

**B**

| Treatment | N | Nectar volume (µl) | | |
|---|---|---|---|---|
| | | mean | SE | P (Wilcoxon rank-sum) |
| High | 49 | 0.33 | 0.04 | |
| High in jar | 59 | 0.38 | 0.05 | 0.45 |
| Low in jar | 58 | 0.38 | 0.05 | 0.79 |
| Int. | 51 | 0.32 | 0.03 | 0.88 |
| Low | 81 | 0.46 | 0.06 | 0.36 |

**Table S2. Experiment 2**. A. **Mean sugar concentration** differed significantly between flowers exposed to “High” frequencies, compared to flowers exposed to “Low" frequencies (P=0.0002), but did not differ significantly between flowers exposed to “High” frequencies, compared to flowers exposed to "Low in jar", "High in jar" or "Intermediate" frequencies (P>0.12) even when not adjusting for multiple comparisons (thus obtaining low, non-conservative, p-values shown in brackets). Mean sugar concentration did not differ significantly between flowers exposed to “High in Jar” condition, compared to flowers exposed to "Low in jar" condition (P=0.64, not in table). Mean sugar concentration ratio differed significantly between flowers exposed to “High” frequencies, compared to flowers exposed to “Low” frequencies (P=0.002), but did not differ significantly between flowers exposed to “High” frequencies, compared to flowers exposed to "Low in jar", "High in jar" or "Intermediate" frequencies (P>0.20). Mean sugar concentration ratio did not differ significantly between flowers exposed to “High in jar” condition, compared to flowers exposed to "Low in jar" condition (P=0.86, not in table). B. **Mean nectar volume** did not differ significantly between any of the groups (P≥0.36).

| Treatment | N | Sugar concentration (%) | | | Sugar concentration ratio | | | Nectar volume (µl) | | |
|---|---|---|---|---|---|---|---|---|---|---|
| | | **mean** | **SE** | **P (Wilcoxon rank-sum)** | **mean** | **SE** | **P (Wilcoxon rank-sum)** | **mean** | **SE** | **P (Wilcoxon rank-sum)** |
| Low | 55 | 15.85 | 0.63 | 0.008 | 0.71 | 0.04 | 0.12 | 0.27 | 0.04 | 0.6 |
| High | 57 | 13.64 | 0.37 | | 0.62 | 0.02 | | 0.27 | 0.03 | |

**Table S3. Experiment 3**. Mean sugar concentration differed significantly between flowers exposed to “Low” frequencies and “High” frequencies (P=0.008). Mean sugar concentration ratio (P=0.12) and Mean nectar volume did not differ significantly between the groups (P=0.6).

| Experiment | Sugar concentration (%) | | Nectar volume (μl) | |
|---|---|---|---|---|
| | mean | SE | mean | SE |
| Outdoor – summer 2014 | 26.7 | 0.62 | 27.26 | 1.5 |
| Indoor – summer 2015 | 25.16 | 0.36 | 8.81 | 0.39 |
| Indoor – fall 2016 | 20.85 | 0.3 | 7.73 | 0.29 |
| Indoor – spring 2016 | 23.06 | 0.48 | 6.72 | 0.3 |

**Table S4. Nectar characteristics before sound treatment – all experiments.** Mean sugar concentration before treatments was significantly higher ($p<10^{-4}$) in experiments performed in summer than in the spring, and was higher ($p<10^{-4}$) in the spring than in the fall. Thus, the absolute value of sugar concentration could not be compared across seasons. Nectar volume was dramatically higher in the outdoor experiment (where the plants were grown outdoors, but tested indoors) than in any of the other experiments.
